# High throughput single-cell genome sequencing gives insights into the generation and evolution of mosaic aneuploidy in *Leishmania donovani*

**DOI:** 10.1101/2021.05.11.443577

**Authors:** Gabriel H. Negreira, Pieter Monsieurs, Hideo Imamura, Ilse Maes, Nada Kuk, Akila Yagoubat, Frederik Van den Broeck, Yvon Sterkers, Jean-Claude Dujardin, Malgorzata A. Domagalska

## Abstract

*Leishmania*, a unicellular eukaryotic parasite, is a unique model for aneuploidy and cellular heterogeneity, along with their potential role in adaptation to environmental stresses. Somy variation within clonal populations was previously explored in a small subset of chromosomes using fluorescence hybridization methods. This phenomenon, termed mosaic aneuploidy (MA), might have important evolutionary and functional implications but remains under-explored due to technological limitations. Here, we applied and validated a high throughput single-cell genome sequencing method to study for the first time the extent and dynamics of whole karyotype heterogeneity in two *Leishmania* clonal populations representing different stages of MA evolution in vitro. We found that drastic changes in karyotypes quickly emerge in a population stemming from an almost euploid founder cell. This possibly involves polyploidization/hybridization at an early stage of population expansion, followed by assorted ploidy reduction. During further stages of expansion, MA increases by moderate and gradual karyotypic alterations. MA usually affected a defined subset of chromosomes, of which some display an enrichment in snoRNA genes which could represent an adaptative benefit to the amplification of these chromosomes. Our data provide the first complete characterization of MA in *Leishmania* and pave the way for further functional studies.

**Note to the BioRxiv community:** The present preprint is a revision of an older preprint posted on 06th March 2020 on BioRxiv (https://www.biorxiv.org/content/10.1101/2020.03.05.976233v1). Here we included two extra samples in our single-cell genome sequencing (SCGS) analysis – the BPK081 cl8 clone (a nearly euploid strain) and a population consisting of a mixture of four *L. donovani* strains which was used as control for high levels of mosaicism in aneuploidy and for estimation of doublets. We also upgraded the bioinformatics pipeline to determine single-cell karyotypes and performed new fluorescence in situ hybridization (FISH) analysis. The new findings observed especially in the BPK081 cl8 led to a reformulation of the text, a new hypothesis for the evolution of mosaicism and a general restructuring of the article. Therefore, the older preprint is obsolete.

## Introduction

Aneuploidy, i.e., an imbalance in the copy number of chromosomes in a cell, occurs in a wide range of organisms, including both non- and pathogenic unicellular eukaryotes, such as *Saccharomyces cerevisiae, Candida albicans, Cryptococcus neoformans* and *Leishmania* spp, but also in different types of human cancer cells (Downing et al., 2011; Holland and Cleveland, 2009; Mulla et al., 2014; Rogers et al., 2011; Selmecki et al., 2006; Sterkers et al., 2011). Although generally considered to be detrimental in multicellular organisms, aneuploidy can also be beneficial, in particular for unicellular organisms facing drastic changes in the environment (Gilchrist and Stelkens, 2019; Siegel and Amon, 2012). In pathogens, aneuploidy facilitates rapid adaptation to environmental stresses through changes in gene dosage and may have an impact on both virulence and the development of drug resistance (Beach et al., 2017; Gerstein et al., 2015; Gilchrist and Stelkens, 2019; Hirakawa et al., 2017; Hu et al., 2011; Ni et al., 2013; Reis-Cunha et al., 2017).

*Leishmania*, a genus of digenetic protozoan parasites, is emerging as a unique model for aneuploidy (Mannaert et al., 2012). These parasites are responsible for a spectrum of clinical forms of leishmaniasis worldwide and cause 300,000 new cases per year (WHO, 2020). They can be found in two forms during their life cycle: as an extracellular promastigote in the midgut of phlebotomine sand fly vectors and exclusively as intracellular amastigote inside mammalian host phagocytic cells. Thus, *Leishmania* parasites are adapted to these two drastically different environments. From a molecular point of view, *Leishmania*, as other trypanosomatids, is unique in the Eukaryota domain (Adl et al., 2012). This includes the genomic organization in long polycistronic units, the near absence of transcription initiation regulation by RNA polymerase II promoters with gene expression regulation almost exclusively through post-transcriptional mechanisms, and its remarkable genomic plasticity (Clayton, 2019; Reis-Cunha et al., 2017). The *Leishmania* genome is generally considered to be diploid, although all *Leishmania* genomes analyzed hitherto display aneuploidy afecting at least one chromosome, i.e., a polysomy in Chr31. Moreover, high levels of ‘average’ aneuploidy (average will be used throughout this paper for features derived from bulk analyses of population of cells) affecting other chromosomes are commonly found by bulk genome sequencing (BGS) of in vitro cultured promastigotes (Downing et al., 2011; Rogers et al., 2011). This average aneuploidy is highly dynamic and changes when cultivated parasite populations are exposed to different environments such as the vector, the vertebrate host or in response to drug pressure (Dumetz et al., 2017; Shaw et al., 2016; Ubeda et al., 2008). In fact, changes in average aneuploidy pattern and not variation in nucleotide sequence are the first genomic modifications observed at populational level during the course of experimental selection of drug resistance (Dumetz et al., 2018; Shaw et al., 2016). Given that these alterations in average somies are reflected in the average amount of corresponding transcripts, and to a certain degree, of proteins, it has been proposed that aneuploidy allows *Leishmania* to adapt by means of rapid changes in gene dosage (Barja et al., 2017; Cuypers, 2018; Dumetz et al., 2017).

*Leishmania* parasites exhibit a remarkable cellular heterogeneity in the form of mosaic aneuploidy, where individual daughter cells originating from a single parent (i.e., a clonal population) may display distinct somies (Lachaud et al., 2014; Sterkers et al., 2011). The full extent of mosaic aneuploidy in *Leishmania* and its dynamics during adaptation to new environment remains largely unexplored due to technological limitations. The only estimation of karyotype heterogeneity was based on the FISH studies of a small set of chromosomes, where it was speculated that thousands of karyotypes may co-exist in a clonal population of *Leishmania* promastigotes (Sterkers et al., 2011). Mosaicism was proposed to provide a source of functional diversity within a population of *Leishmania* cells, through gene dosage, but also through changes in heterozygosity (Barja et al., 2017; Sterkers et al., 2012). This diversity of karyotypes would provide an adaptive potential to unpredictable environmental changes during the parasite’s life cycle or drug pressure caused by patient treatment (Barja et al., 2017; Sterkers et al., 2012).

Here, we applied and validated for the first time a high throughput, droplet-based platform for single cell genome sequencing (SCGS) of thousands of individual *Leishmania* promastigotes. This allowed the assessment of the degree and the dynamics of the evolution of mosaic aneuploidy in two clonal populations in vitro representing different stages of adaptation to culture conditions. Based on our study, we propose that the early stages of adaptation are characterized by rapid and drastic changes in karyotypes, allowing initial establishment of highly aneuploid cells in a population of almost euploid parasites. In the next steps, the existing highly aneuploid karyotypes further evolve through gradual and moderate changes in somies resulting in a population of aneuploid cells displaying closely related karyotypes. Our findings strongly support the hypothesis that mosaic aneuploidy is a constitutive feature of *Leishmania* parasites, representing a unique source of functional diversity.

## Materials and Methods

### Parasites

In the present paper we use the terms population, strain and clone as defined in the supplementary text. *L. donovani* promastigotes were maintained at 26 °C in HOMEM medium (Gibco, ThermoFisher) supplemented with 20% Fetal Bovine Serum, with regular passages done every 7 days at 1/25 dilutions. The clones BPK282 cl4 and BPK081 cl8 were derived from two strains adapted to culture: MHOM/NP/02/BPK282/0 and MHOM/NP/02/BPK081/0 (Imamura et al., 2016). These clones were submitted to SCGS at 21 (∼126 generations) and 7 passages (∼56 generations) after cloning respectively (supp. fig.1). Four strains were mixed to create an artificial ‘super-mosaic’ population of cells (further called super-mosaic): BPK475 (MHOM/NP/09/BPK475/9), BPK498 (MHOM/NP/09/BPK498/0), BPK506 (MHOM/NP/09/BPK506/0) and HU3 (MHOM/ET/67/HU3). They were kept in vitro for several passages after isolation from patients (respectively 41, 60, 47 and more than 24) and mixed at equivalent ratio just before preparation for SCGS.

### Single-cell suspensions preparation and sequencing

Promastigotes at early stationary phase (day 5) were harvested by centrifugation at 1000 rcf for 5 min, washed twice with PBS 1X (calcium and magnesium-free) + 0.04% BSA, diluted to 5×10^6^ parasites/mL and passed through a 5 µm strainer to remove clumps of cells. After straining, volume was adjusted with PBS 1X + 0.04% BSA to achieve a final concentration of 3×10^6^ parasites/mL. The absence of remaining cell doublets or clumps in the cell suspension was confirmed by microscopy. Cell viability was estimated by flow cytometry (BD FACSverse^TM^) using the NucRed^TM^ Dead 647 probe (Life technologies^TM^) following the recommendations of the manufacturer and in all samples was estimated as higher than 95%. SCGS was performed using the Chromium^TM^ single-cell CNV solution (scCNV) from 10X Genomics^TM^. To target an average of 2000 sequenced cells per sample, 4.2 µL of the cell suspensions were used as input, and cell encapsulation, barcoding, whole genome amplification and library preparation were performed following manufacturer’s recommendations. Sequencing of the libraries was done with an Illumina NovaSeq^TM^ SP platform with 2×150 bp reads.

### Single-Cell Somy estimation

Details about the bioinformatics analysis for somy values determination are provided in the supplementary material. In summary, sequence reads were associated to each sequenced cell based on their barcodes and mapped to a customized version of the reference *L. donovani* genome LdBPKv2 (Dumetz et al., 2017) using the Cell Ranger DNA^TM^ software (10X Genomics). The matrix generated by the software with the number of mapped reads per 20kb bins was used as input to a custom script written in R (R Core Team, 2013). In this script, bins with outlier values were excluded, and the mean normalized read depth (MNRD) of each chromosome was calculated for each cell. Cells displaying a high intra-chromosomal variation were removed from downstream analysis. In order to establish the baseline ploidy of each cell, the MNRD values were multiplied by the scale factor (Sc), defined for each cell as the lowest number between 1.8 and 5 that leads to the shortest distance to integers when all MNRD values are multiplied by it. The MNRD values multiplied by Sc are referred here as ‘raw somies’. To convert the raw somies (continuous) into integer copy numbers (discrete), a univariate gaussian mixture-model was built for each chromosome by an expectation-maximization algorithm based on the distribution of the raw somy values between all cells of the same sample using the Mixtools package (Benaglia et al., 2009). For each possible integer somy, a gaussian mixture-model was generated and each raw somy value was assigned to the rounded mean of the gaussian to which it has higher probability of belonging to.

### Karyotype identification and network analysis

A karyotype was defined as the combination of integer somies of all chromosomes in a cell. Karyotypes were numerically named according to their frequency in the sequenced population. To generate the network representing the dissimilarities between the karyotypes, a pairwise distance matrix was built based on the number of different chromosomes between all karyotypes in a sample, and used to create a randomized minimum spanning tree with 100 randomizations, using the Pegas R package (Paradis, 2018, 2010). The network visualization was made with the visNetwork package (Almende B.V. et al., 2019).

### Doublet detection

The relative fraction of doublets within the super mosaic population has been estimated based on the high number of SNPs found in the HU3 strain when compared to the *L. donovani* reference genome. The three other strains in the super mosaic only show a limited number of SNPs in contrast. Potential doublets were identified by looking for mixture of both SNP profiles (HU3 and non-HU3) in assumed single-cell data. This approach was applied using an in-house developed algorithm and the Demuxlet algorithm (Kang et al., 2018), both approaches leading to identical results (see Supplementary Text).

### DNA probes and fluorescence in situ hybridization

DNA probes were either cosmid (L549 specific of chromosome 1) or BAC (LB00822 and LB00273 for chromosomes 5 and 22 respectively) clones that were kindly provided by Peter Myler (Seattle Biomedical Research Institute) and Christiane Hertz-Fowler (Sanger Centre). DNA was prepared using Qiagen Large-Construct Kit and labelled with tetramethyl-rhodamine-5-dUTP (Roche Applied Sciences) by using the Nick Translation Mix (Roche Applied Sciences) according to manufacturer instructions. *Leishmania* cells were fixed in 4% paraformaldehyde then air-dried on microscope immunofluorescence slides, dehydrated in serial ethanol baths (50–100%) and incubated in NP40 0.1 % for 5 min at RT. Around 100 ng of labelled DNA probe was diluted in hybridization solution containing 50% formamide, 10% dextran sulfate, 2× SSPE, 250 µg.mL^−1^ salmon sperm DNA. Slides were hybridized with a heat-denatured DNA probe under a sealed rubber frame at 94 °C for 2 min and then overnight at 37 °C and sequentially washed in 50% formamide/2× SSC at 37 °C for 30 min, 2× SSC at 50 °C for 10 min, 2× SSC at 60 °C for 10 min, 4× SSC at room temperature. Finally, slides were mounted in Vectashield (Vector Laboratories) with DAPI. Fluorescence was visualized using appropriate filters on a Zeiss Axioplan 2 microscope with a 100× objective. Digital images were captured using a Photometrics CoolSnap CCD camera (Roper Scientific) and processed with MetaView (Universal Imaging). Z-Stack image acquisitions (15 planes of 0.25 µm) were systematically performed for each cell analyzed using a Piezo controller, allowing to view the nucleus in all planes and to count the total number of labelled chromosomes. Around 200 cells [187-228] were analyzed per chromosome.

### Bulk Genome Sequencing (BGS)

Genomic DNA from the BPK282 cl4 and BPK081 cl8 clones was extracted in bulk using the QIAmp^TM^ DNA Mini kit (Qiagen) following manufacturer’s recommendations. PCR-free whole genome sequencing was performed on the Illumina NovaSeq platform using 2×150 bp paired reads. Reads are mapped to the reference genome *L. donovani* LdBPKv2 (available at ftp://ftp.sanger.ac.uk/pub/project/pathogens/Leishmania/donovani/LdBPKPAC2016beta/) using BWA (version 0.7.17) with seed length set to 100 (Li and Durbin, 2009). Only properly paired reads with a mapping quality higher than 30 were selected using SAMtools (Li et al., 2009). Duplicates reads were removed using the RemoveDuplicates command in the Picard software (http://broadinstitute.github.io/picard/). The average somy values were calculated as described previously (Downing et al., 2011), by dividing the median sequencing depth of a chromosome by the overall median sequencing depth over all chromosomes, and multiplying this ratio by 2. These values were used to define an average karyotype for the sequenced population of cells (Kp).

### Gene Ontology analysis and in silico screening for small RNA

Gene Ontology (GO) classes were obtained from TriTrypDB release 49 (Aslett et al., 2009). As the genome sequence stored on TriTrypDB does not correspond with the reference genome used in this work, the GO annotation was obtained by mapping back all genes to our reference genome using BlastP (Altschul et al., 1997). Clustering of the different chromosomes based on their assigned GO classes was performed using the prcomp command in R.

The Rfam (Kalvari et al., 2021) database version 14.4 was used to screen the *L. donovani* BPK282 reference genome using the cmscan algorithm as implemented in Infernal (Nawrocki and Eddy, 2013) using default parameters and setting the search space parameter to 64.

## Results

### High throughput single-cell genome sequencing as a reliable tool to explore karyotype heterogeneity in *Leishmania* populations

We applied high throughput single-cell genome sequencing (SCGS) to address mosaic aneuploidy in promastigotes of two *Leishmania* clones differing substantially in average aneuploidy (refered here as the ‘average populational karyotype’, or Kp) as revealed by Bulk Genome Sequencing (BGS): (i) BPK282 cl4, an aneuploid clone showing 7 chromosomes with an average trisomy apart from the usual average tetrasomy in Chr31 and (ii) BPK081 cl8, showing an average disomy for all chromosomes except Chr31 (average tetrasomic); for simplicity, we will call BPK081 cl8 the ‘diploid’ clone. First analyses of the SCGS data were made with the Cell Ranger DNA^TM^ pipeline. Although the software was developed for mammalian genomes, which are up to 2 orders of magnitude larger than *Leishmania*’s nuclear genome, it allowed detecting (i) aneuploidy, (ii) mosaicism and (iii) large intrachromosomal CNVs, as, for instance, the H- and M-amplicons (Downing et al., 2011) in Chr23 and Chr36 respectively (Suppl. fig 2). However, technical artifacts were noticed especially in BPK081 cl8, where the software’s GC bias correction algorithm, designed for the mammalian genome which display a lower average GC content compared to *Leishmania*, ended up overcompensating the depth of bins with high GC content (Suppl. fig 2). Because of that and given our main goal of using SCGS to study mosaic aneuploidy, we built our own analytical bioinformatic pipeline with a higher emphasis on estimating whole chromosomes copy numbers rather than local CNVs (Suppl. fig 3).

We evaluated the SCGS method and our analytical pipeline by first addressing their ability to explore karyotype heterogeneity among *Leishmania* cells of clones BPK282 cl4 and BPK081 cl8. Using our analytical pipeline, we identified 208 different karyotypes among the 1516 filtered cells of BPK282 cl4 and 117 karyotypes among the 2378 filtered cells of BPK081 cl8 (fig.1 A-B, Suppl. fig 5 A-B). Moreover, the cumulative SCGS profile of each clone was consistent with their respective Kp (fig. 1A and 1B, left panel). Notably, Chr13, which displays a non-integer average somy value (2.26) in the Kp of BPK282 cl4, was found as disomic and trisomic at relatively high proportions in the SCGS, resulting in a similar cumulative somy (2.34). As expected, the vast majority of cells in BPK081 cl8 displayed an almost diploid karyotype, with only Chr31 displaying a tetrasomy as expected. Small subpopulations of cells displaying highly aneuploid karyotypes were also observed in BPK081 cl8 (discussed below).

**Figure 1.**
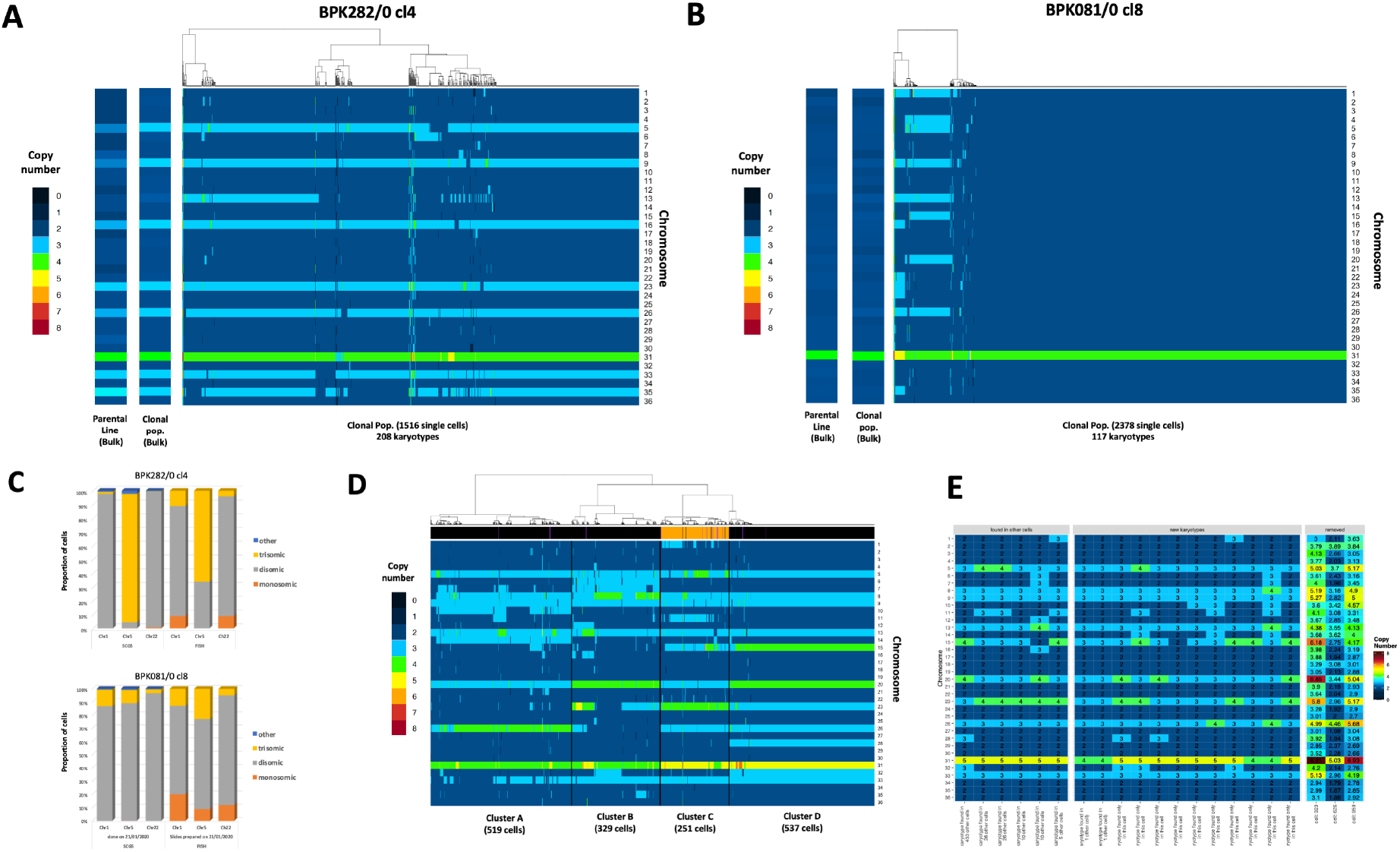
Mosaic aneuploidy in BPK282 cl4 and BPK081 cl8 clones revealed by SCGS and validation of the method. **A-B.** Heat maps displaying the copy number of all 36 chromosomes of promastigotes from BPK282 cl4 (A) or BPK081 cl8 (B) clones (main panels). Each column represents a single parasite. The number of sequenced promastigotes and karyotypes found in each sample is described in the x axis. In each panel, two insets display the Kp of the clonal population used in the SCGS and their respective parental strain. **C.** Comparison between FISH and SCGS. The proportion of cells displaying monosomy, disomy or trisomy for chromosomes 1, 5 and 22 in each method is represented. **D.** Heat map displaying the karyotypes of the promastigotes from 4 different strains mixed in a single SCGS run. Cells were hierarchically clustered according to their karyotypes, forming 4 major clusters. The number of cells in each cluster is indicated in the x axis. The bar at the top of the heatmap indicate if the SNP profile of the cell correspond to a BPK strain (black), a HU3 strain (orange) or a doublet (purple). **E.** Karyotypes of cells marked as doublets. The number of other cells displaying the same karyotype as the doublet is indicated in the x axis labels. Cells that were removed from analysis due to high intra-chromosomal variation and therefore did not have their somy values converted to integers are separated in the right panel, displaying their raw somy values instead. The integer somy values (left panels) or the raw somy values (right panel) are numerically indicated inside

Mosaic aneuploidy in *Leishmania* has been studied so far with fluorescence in situ hybridization (FISH), the only alternative method available hitherto to estimate the copy number of some chromosomes in individual *Leishmania* cells. As a mutual benchmark of both FISH and SCGS methods, we submitted cells from both BPK282 cl4 and BPK081 cl8 to FISH to estimate the copy number of chromosomes 1, 5 and 22 and to compare the obtained results with the values observed in our SCGS data (fig. 1C). Overall, for each chromosome, the same predominant somy was observed with both methods, even when the predominant somy was different between clones. For instance, FISH and SCGS report Chr5 in BPK282 cl4 as trisomic in most cells, while it is reported as mainly disomic in BPK081 cl8 also by both techniques. Most discrepancies between the proportions obtained by both methods are within the 10% error margin previously estimated for FISH (Sterkers et al., 2011 and unpublished results). The main exception is Chr5 in BPK282, which is estimated as trisomic in 93% of the cells with SCGS and 66% with FISH. However, SCGS reports proportions which are more consistent with the average somy values obtained by the BGS analysis of each clone. For instance, the weighted mean between somy values obtained with SCGS for Chr5 in BPK282 cl4 results in an average somy of 2.95, which is very similar to the average somy value obtained by BGS (2.97), whereas with FISH, the average somy is lower (2.66), suggesting that the proportions observed with SCGS are more accurate.

We executed an extra experiment to evaluate the performance of SCGS in dealing with populations with highly heterogeneous karyotypes. In this experiment, a ‘super-mosaic’ population was generated by mixing 4 different *L. donovani* strains that display very distinct Kp’s(Imamura et al., 2016), into a single SCGS run. A total of 1900 promastigotes were individually sequenced, of which, 1636 remained after data filtering. This ‘super mosaic’ population displayed a high aneuploidy diversity: 388 identified karyotypes in total. As expected, the 1636 promastigotes formed four distinct clusters based on their integer somy values, with discrete differences in the aneuploidy patterns between each cluster (fig. 1D). Since one of the strains (HU3) used in this super mosaic is phylogenetically distant from the other 3 strains (BPK475, BPK498 and BPK506), we could distinguish HU3 promastigotes from the others based on their SNP profiles. Interestingly, all HU3 cells were grouped together in cluster C (fig. 1D – orange lines in the annotation bar), suggesting that the discrete karyotypic differences between the major clusters reflect differences among the aneuploidy profiles of the four strains, so that each cluster likely represents one of the strains. Thus, this experiment demonstrates that SCGS is effective in distinguishing karyotypes even in very complex populations.

The ‘super-mosaic’ population was also used to estimate the frequency of doublets, i.e, the inclusion in a single droplet of two or more cells sharing the same 10X barcode. Based on the SNP profile of the HU3 line, each dataset with the same barcode containing either none of the HU3-specific SNPs (< 5% of the SNPs), or almost all of the HU3-specific SNPs (> 95% of the SNPs) were defined as singlets, while doublets contained a mixture of HU3-speficic SNPs and positions resembling the reference genome. Using this approach, from the 293 cells that were predicted as HU3 based on their SNP profile (including cells removed from karyotype estimation), 21 were predicted as doublets (fig. 1D; purple lines in the annotation bar), with a detection rate of SNPs varying between 14% and 58%. Since doublets formed by two HU3 cells would still be defined as a singlet and given that HU3 cells correspond to 15,4% of the population, we assumed that the 21 detected HU3+BPK doublets correspond to 84,6% of the total number of doublets containing an HU3 cell. Thus, we estimate that there are ∼4 (the extra 15,4%) additional HU3+HU3 doublets, resulting in a total of 25 doublets. Extrapolating this fraction of 25 out of 293 HU3 cells to the whole single-cell population would correspond to a relative fraction of doublets of 8,53%, a frequency which is higher than anticipated for mammal cells according to the manufacturer’s guidelines (∼1,4%). From the 21 detected doublets, 3 were originally removed from karyotype estimation due to high intra-chromosomic variation, and 6 displayed a karyotype that was also found in other cells. However, 11 karyotypes were exclusively found in one of the detected doublets (fig. 1E), indicating that a fraction of the low-occurrence karyotypes might be artifacts due to doublets.

### BPK282 and BPK081 cells display different patterns of karyotype evolution during clonal expansion

After validating the SCGS method for resolving complex karyotype heterogeneity in *Leishmania*, we returned to the data of BPK282 cl4 and BPK081 cl8 to characterize the karyotypes that are present in each clone. In BPK282 cl4, the most frequent karyotypes were very similar to each other, diverging by copy number changes in 1 to 3 chromosomes when compared to the most frequent karyotype (kar1 – fig. 2A). In BPK081 cl8, however, the nearly diploid kar1, which was present in 82% of the cells, and the 2 next most abundant karyotypes showed very different aneuploidy profiles, diverging by copy numbers of 8 to 10 chromosomes (fig. 2B). In addition, in both clones, the most frequent karyotype (kar1) is similar to the Kp of the respective parent strain from which each clone was derived (fig. 1 A-B, left panel), suggesting that, in each clone, kar1 corresponds to the karyotype of the founder cell, and thus, the other karyotypes of each population arose from their respective kar1.

**Figure 2.**
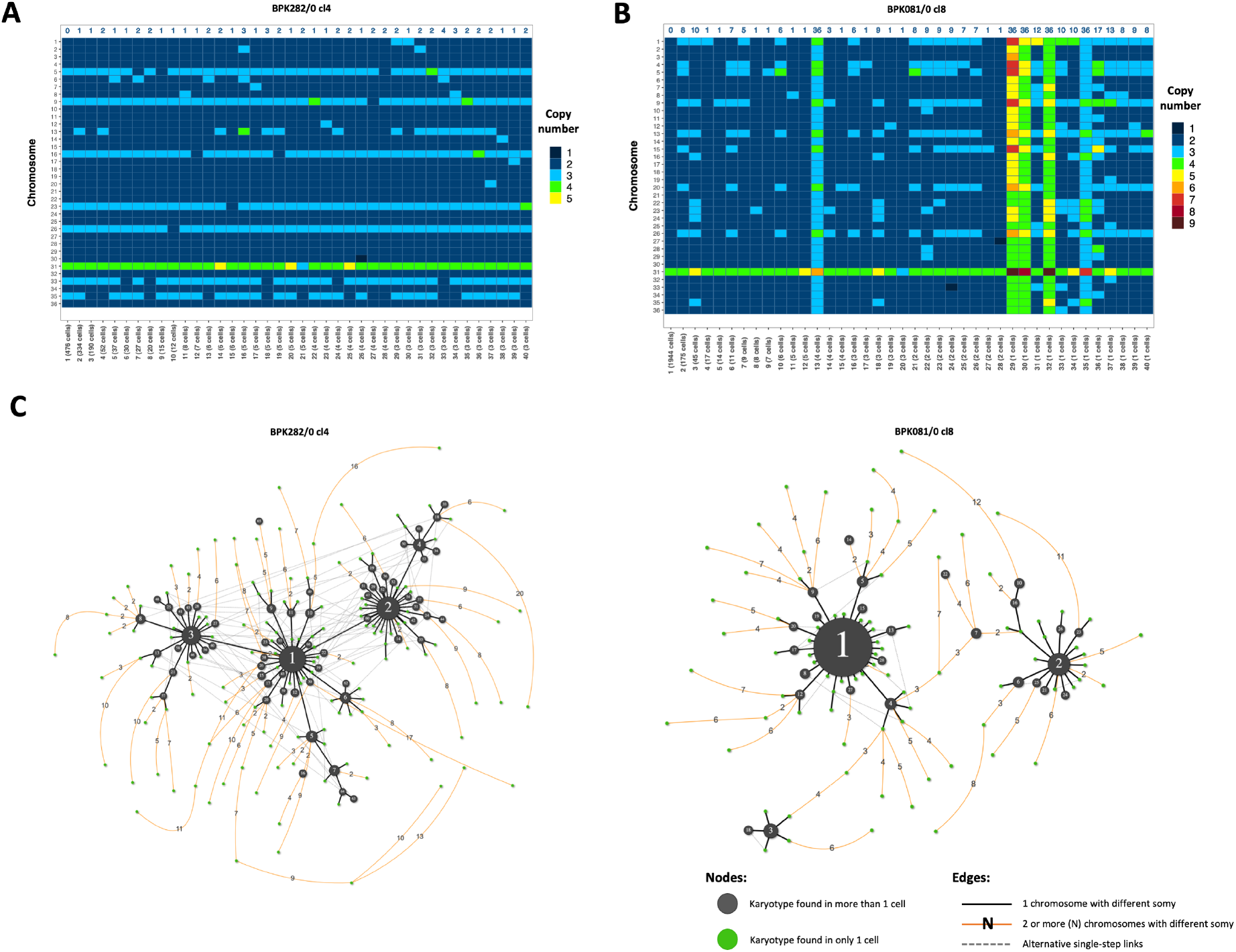
BPK282 cl4 and BPK081 cl8 display different profiles in the dissimilarity relationship between karyotypes. **A-B.** Heat map depicting the 40 most frequent karyotypes in BPK282 cl4 (A) and BPK081 cl8 (B) clones. The blue numbers in the top indicate the total number of chromosomes with a different somy compared to kar1. **C.** Network representing the dissimilarity relationship between karyotypes in each clone. Black nodes represent karyotypes found in more than one cell, with their size proportional to the number of cells. Green nodes indicate karyotypes which occur only once. Black lines link two karyotypes which diverge by a somy difference in a single chromosome, while orange lines link karyotypes diverging by two or more chromosomes with different somy, with the number of divergent chromosomes indicated in the edge. Dashed grey lines show alternative links between karyotypes with a single somy divergency. Polyploidy karyotypes were not included

To develop a hypothesis of the karyotype evolution during expansion of both BPK282 cl4 and BPK081 cl8 populations, we built a dissimilarity network based on the number of chromosomes with different copy numbers between each karyotype found in each population (fig. 2C). Both populations of cells are at different stages of expansion (about 126 and 56 generations since cloning, respectively), but we observe in each of them a proportionally comparable number of somy changes events (steps in the network): (i) for BPK282 cl4, 514 steps/126 generations/1516 sequenced cells = 0.0027 and (ii) for BPK081cl8, 260 steps/56 generations/2378 sequenced cells = 0.002. However, distinct patterns are observed between both clones. In BPK282 cl4, the most frequent karyotypes (black nodes) are linked to each other by somy changes in only single chromosomes (black lines). Assuming kar1 as the founder of this population, almost every frequent karyotype can be traced back to it through cumulative single copy number alterations. In contrast, the network of BPK081 cl8 shows a very distinct pattern (fig. 2C). Here, the 3 most frequent karyotypes are distant from one another and lack single-step intermediates between them.

### Selective forces restrict high frequencies of polysomies to a specific group of chromosomes

We and others have demonstrated that high frequencies of polysomies were restricted to a specific subset of chromosomes when comparing the Kp’s of 204 *L. donovani* strains previously analyzed by BGS (Barja et al., 2017; Imamura et al., 2016). To address if the same applies to single *Leishmania* cells, we created a diverse artificial population by randomly selecting and merging the data of equal numbers of single cells from BPK282 cl4 and BPK081 cl8 as well as from each cluster of the super mosaic, assuming each cluster represents one of the mixed strains. In this artificial population, we observed that at least 16 chromosomes are consistently disomic in the vast majority of cells in a clone/strain-independent manner (fig. 3A). All these chromosomes also show an average disomy in the Kp of most of the 204 strains mentioned above (supp. fig. 7A-B). Conversely, apart from the usually tetrasomic Chr31, 8 chromosomes (Chr5, Chr8, Chr9, Chr13, Chr20, Chr23, Chr26 and Chr33) are found with 3 or more copies in most cells of BPK282 cl4 and BPK081 cl8, again fitting with previous observations made on the 204 *L. donovani* strains (Barja et al., 2017; Imamura et al., 2016). However, it is unclear whether (i) the disparity in the frequency of polysomies between chromosomes is due to intrinsic differences in the chances of overamplification of each chromosome along the expansion of the population (some chromosomes being specifically ‘unstable’) or (ii) if every chromosome has the potential to become polysomic but the expansion of polysomies in a population is determined by selective pressures. To address this, we revisited the karyotype network of each population (including the ‘super-mosaic’ – supp. fig. 7C), to investigate which were the chromosomes that were more prone to somy alterations in the rare karyotypes (i.e., karyotypes occurring in only a single cell), compared to the common karyotypes (i.e., karyotypes occurring in 2 or more cells) (fig. 3B). As expected, the 16 chromosomes which are predominantly found as disomic display little, if any, alteration events in their copy numbers in the common karyotypes. However, between the rare karyotypes, all chromosomes are susceptible to somy alterations with relatively similar frequencies, although polysomy-prone chromosomes still display a higher alteration frequency (p-value < 0.0001 – supp. fig. 7D). These observations suggest that the capacity for aneuploidy is not restricted to a specific group of ‘unstable’ chromosomes.

**Figure 3.**
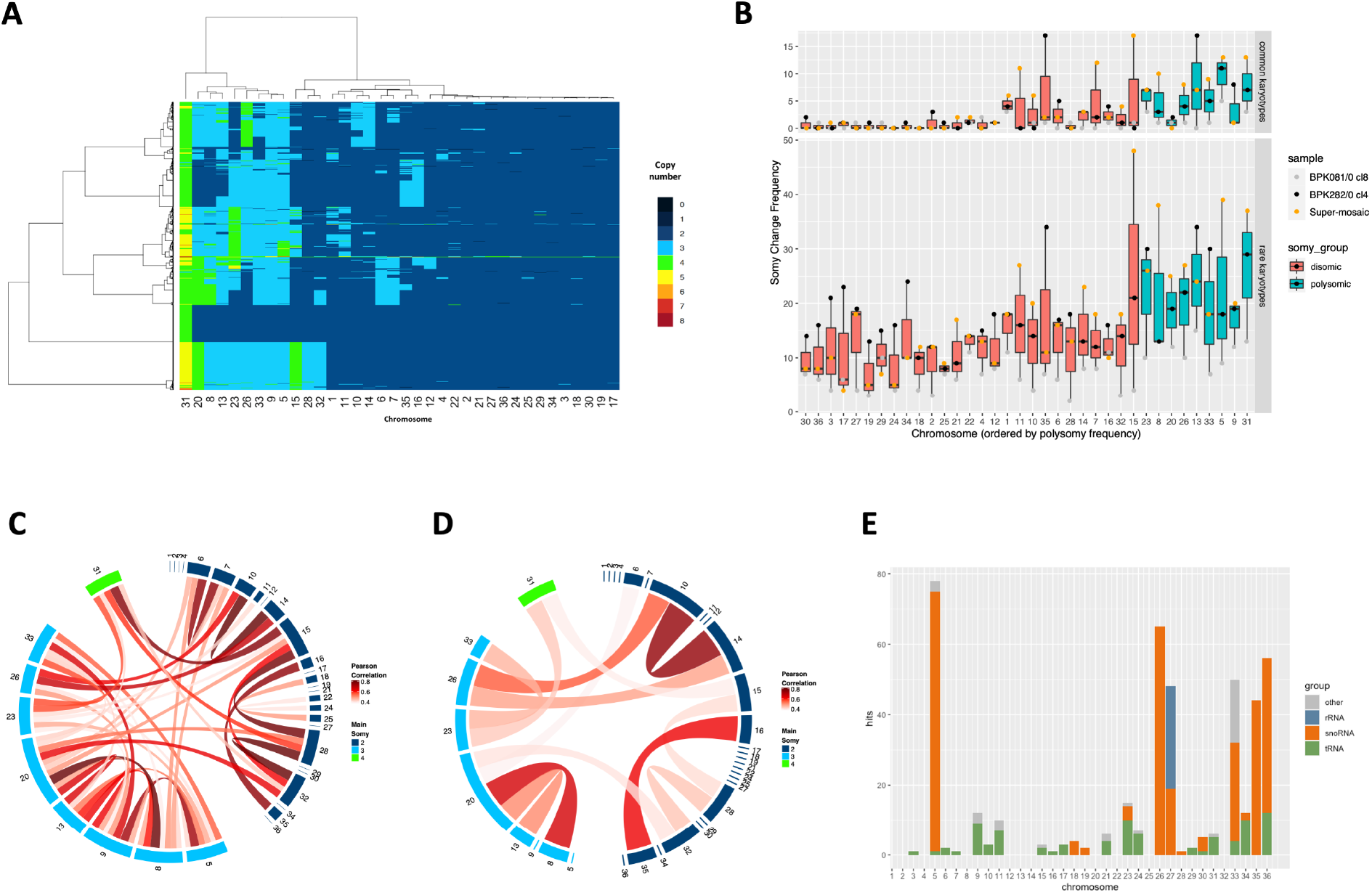
High frequencies of polysomies are restricted to a group of chromosomes. **A.** Heat map depicting the copy number of the 36 chromosomes across promastigotes from different clones/strains. Here, 251 promastigotes of each cluster of the mixed sample and from BPK282 cl4 and BPK081 cl8 are represented. Chromosomes are hierarchically clustered based on their somy values. **B.** Boxplot indicating the number of somy change events for each chromosome among the common karyotypes (found in 2 or more cells – top panel) or the rare karyotypes (found in only one cell - bottom) in the 3 samples submitted to SCGS. **C-D.** Chord diagrams representing the Pearson correlation between the somies of all chromosomes among cells displaying the common karyotypes (C) or the rare karyotypes (D). Only correlations higher than 0.4 and with p.value lower than 0.05 are represented. **E.** Distribution of small non-coding RNAs across *L. donovani* genome. Ribosomal RNAs (rRNA), small nucleolar RNAs (snoRNAs) and transporter RNAs (tRNAs) were identified based on the Rfam database.

We also investigated the role of the synchronous fluctuation in the copy number of multiple chromosomes in determining the abundance of karyotypes. For that, we estimated Pearson correlations between the copy number of chromosomes across equal numbers of cells from all clones/strains sequenced here (supp. fig. 7E). Between the 8 polysomy-prone chromosomes and among the cells with common karyotypes, we observed numerous and relatively strong correlations, with the strongest correlations occurring between Chr5 and Chr9, and Chr8 and Chr20 (fig. 3C). On the other hand, between cells with rare karyotypes, there were fewer and in general weaker correlations (fig. 3D). These observations suggest that the expansion of polysomies in a population happens in an interdependent manner between chromosomes.

### Functional characterization of the polysomy-prone chromosomes

In order to investigate potential features specific to the polysomy-prone chromosomes that could be related to their higher frequency of polysomies, we first applied an unsupervised Gene Ontology (GO) analysis to look for enrichment of biological functions in the polysomy-prone chromosomes. However, no obvious relationships between chromosomal gene content and prevalence of polysomies could be found (supp. fig. 8A). We then tried a supervised approach. Since highly aneuploid karyotypes are more frequently observed in in vitro promastigotes than in amastigotes, we reasoned that the amplification of the polysomic-prone chromosomes might affect pathways related to the promastigote stage. Thus, we selected enriched GO classes which were obtained from a previously published study in which we studied differential expression between promastigote and amastigote cell cultures (Dumetz et al., 2017). The distribution of the corresponding genes on the polysomy-prone chromosomes was compared to the distribution on chromosomes with a stable disomy. However, this approach also did not disclose biological functions located on the amplified chromosomes (supp. fig. 8B). Alternatively to GO analysis, we finally performed an in silico scan for small non-coding RNAs to investigate their distribution throughout the *L. donovani* genome. This suggested an enrichment of small RNAs in some of the polysomy-prone chromosomes, especially small nucleolar RNAs (snoRNAs - fig. 3E). A significant number of hits for snoRNAs are mapped to Chr5, Chr26 and Chr33, which are among the chromosomes with the most frequent polysomies, as well as Chr35, which is trisomic in the majority of BPK282 cl4 cells and is also trisomic in the Kp of several *L. donovani* strains (Imamura et al., 2016). Although preliminary, this observation suggests a potential relationship between the snoRNAs content of a chromosome and its prevalence of polysomies in cultivated promastigotes.

### SCGS reveals particular karyotypes among rare single cells

As shown above, kar2 and kar3 of BPK081 cl8 show a baseline diploidy, i.e., the majority of chromosomes are disomic, with 8 to 10 trisomic chromosomes and tetrasomy or even a pentasomy for for Chr 31. However, we found in the same population 4 cells displaying a karyotype (kar13) with an aneuploidy profile similar to kar2, but with all chromosomes showing one extra copy (two extra copies for Chr31 - fig. 2B); thus in kar11, baseline somies are trisomic, 8 chromosomes (the same as kar2) are tetrasomic and Chr31 is hexasomic, constituting a triploid karyotype (see supplementary text for details on how cells ploidies are determined). Similarly, at least 1 cell showed another karyotype (kar35) with baseline triploidy and aneuploidy on the same chromosomes as kar3 (fig. 2B). Tetraploid karyotypes were also observed among BPK081 cl8 cells, but it is not possible to rule out that these are in fact doublets between two 2n cells with different karyotypes. Noteworthy, tetraploid karyotypes were not found in BPK282 cl4 and the only 3 cells identified with a potential baseline triploidy exhibited an aneuploidy pattern very distinctive from any other karyotype in that population (supp. fig. 6). Moreover, within the BPK282 cl4 and BPK081 cl8 populations, we also observed rare cells displaying chromosomes with an estimated somy of 0 (nullisomy). The bam file of these cells showed that no reads were mapping to these chromosomes, suggesting that in these cells, these chromosomes were absent (fig. 4A). Nullisomic chromosomes were found in all the populations sequenced here: among which, 4 in BPK081 cl8 (0,15% of the sequenced cells) and 15 from BPK282 cl4 (0,88%). Moreover, the aneuploidy profile of these nullisomic cells was not similar to any other karyotype identified in each sample (fig. 4B). Partial chromosome deletions were also observed, as for instance in Chr13 and Chr36 of the cell 688 from BPK282 cl4, in the Chr36 of the cell 266 from BPK081 cl8.

**Figure 4.**
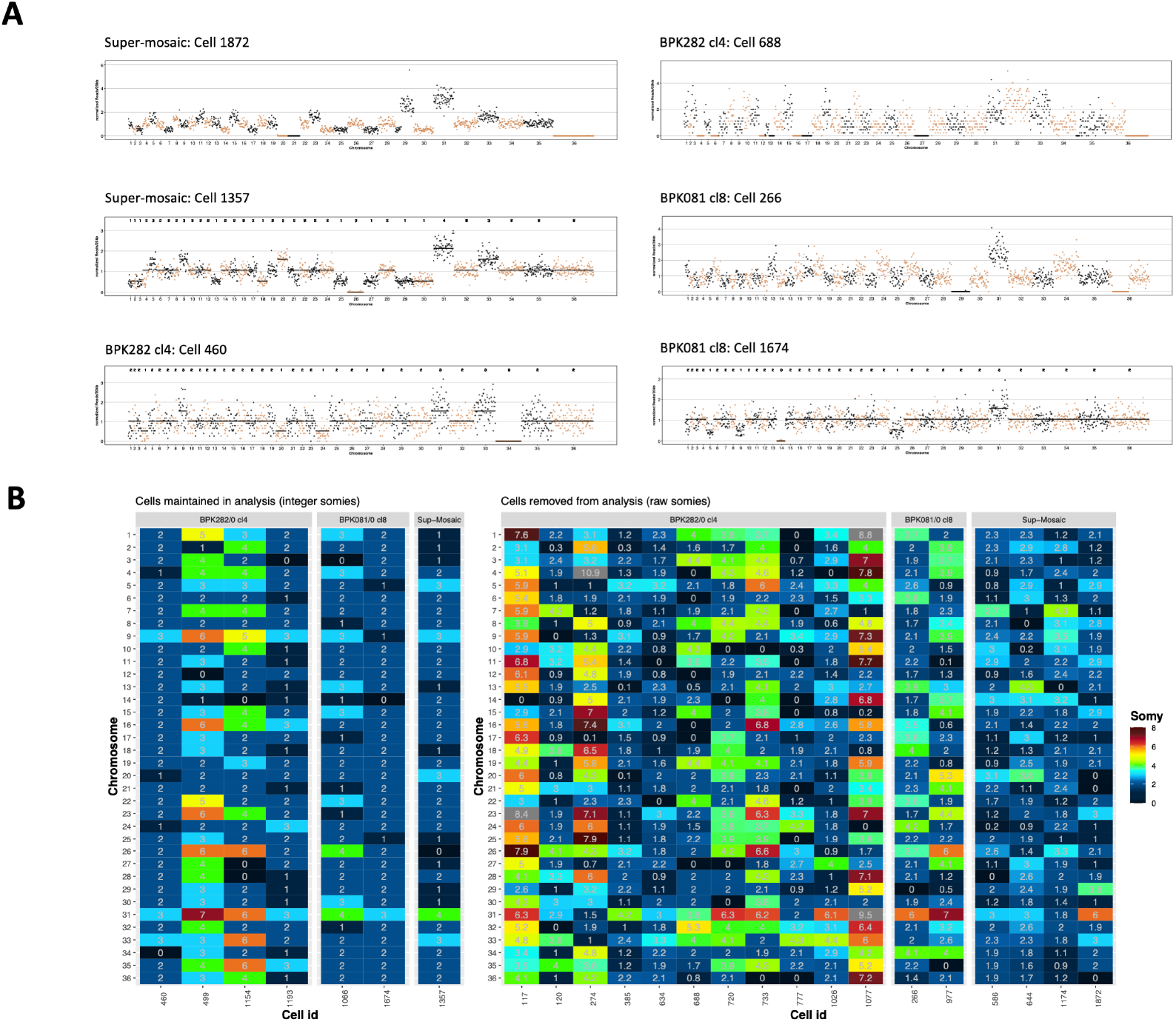
Cells with nullisomic chromosomes. **A.** Example of cells displaying one or more nullisomic chromosomes. The dots represent the normalized read depth of each 20kb bin. The integer somy values calculated for each cell are depicted in the top part of each box for cells that were not excluded from analysis. A black line shows the integer somy values divided by the cell’s scale factor (Sc) for comparison. **B.** Karyotype of all cells with at least one nullisomic chromosome identified in our SCGS data. Cells that were removed from analysis and therefore did not have their somy values converted to integers are separated in the right panel, displaying their raw somy values instead.

## Discussion

Cellular heterogeneity is increasingly implicated as one of the major sources of adaptative potential for unicellular pathogens (Bagamery et al., 2020; Seco-Hidalgo et al., 2015). We explored here a specific manifestation of this phenomenon, i.e., mosaic aneuploidy, in a unique model, *Leishmania*. By applying a high throughput SCGS method, we could determine for the first time the complete karyotype of thousands of individual *Leishmania* cells from two distinct clonal populations in vitro. We found a high level of mosaic aneuploidy, affecting essentially the same, limited subset of chromosomes. We explored the evolution of mosaicism in both populations, starting from two distinct founder karyotypes, one nearly euploid and another highly aneuploid. We highlighted the adaptive potential of mosaic aneuploidy for unicellular organisms such as *Leishmania*, living in rapidly varying environments.

The present SCGS study allowed us to evaluate and extend hypotheses on mosaic aneuploidy in *Leishmania* previously based on FISH measurements (Sterkers et al., 2012, 2011). Although some divergencies were observed here between FISH and SCGS, our data are in agreement with most predictions. Accordingly, mosaic aneuploidy was confirmed in all populations sequenced here, and karyotypes frequency distributions, in particular for BPK282/0 cl4 clone (208 karyotypes among 1516 cells), were similar to the distribution predicted with FISH data obtained for 7 chromosomes of a long-term cultivated *Leishmania major* population (∼250 karyotypes in ∼2000 cells - Sterkers et al., 2012 – fig. 4). In BPK081/0 cl8, proportionally fewer karyotypes were identified compared to BPK282/0 cl4, which might be a consequence of either a reduced tendency of the founder diploid karyotype to somy alterations and/or due to the fact this clone was at an ealier stage of expansion in vitro (∼56 generations, compared to the ∼126 generations in BPK282). Indeed, when normalizing the number of karyotypes, similar values were observed for both clones: respectively 10exp^-4^ and 9exp^-4^ new karyotypes/generation/sequenced cell.

Our SCGS data, however, do not corroborate the previous assumptions that all chromosomes are found with at least two somy states (Sterkers et al., 2012, 2011), as high levels of somy variation were restricted to a subset of chromosomes in our experimental conditions. We also observed a higher tendency of FISH to report trisomies and monosomies in chromosomes which were defined by SCGS as mostly disomic in almost all cells of BPK282/0 cl4 and BPK081/0 cl8 clones, as chr01 and chr22. This discrepancy is likely due to accuracy limitations in FISH.

The SCGS data reported here also allowed us to draw some hypothesis regarding the origin and evolution of mosaic aneuploidy in vitro. We have previously demonstrated that intracellular amastigotes sequenced directly from patient samples usually display a diploid Kp similar to the Kp of the BPK081 cl8 clone, although variations in somies were observed in some samples (Domagalska et al., 2019). However, when these amastigotes were isolated from patients or experimental animals and transformed to promastigotes in vitro, in most cases their Kps progressively evolve towards highly aneuploid profiles (Domagalska et al., 2019; Dumetz et al., 2017; Giovanni Bussotti, a et al., 2018). Thus, the 2 clones here studied provide complementary models to understand the dynamics of the emergence of mosaic aneuploidy in vitro; BPK081/0 cl8 which founder karyotype had the diploid profile, representing an early stage of adaptation to culture; and BPK282/0 cl4, which founder karyotype was already highly aneuploid (likely kar1), representing later stages.

In the BPK081 cl8, a minority of highly aneuploid subpopulations were observed, contrasting with the the founder diploid karyotype (kar1), indicating that at early stages of clonal expansion in culture, the evolution of mosaicism starts with drastic changes in karyotypes, in this case the observed changes in somy of 8 to 10 chromosomes leading to highly aneuploid cells (kar2 and kar3). These drastic changes in somies could occur through cumulative small steps, i.e., somy alterations in single chromosomes at each cell division, followed by fixation and further expansion of the fittest aneuploidies and loss of intermediate links between these karyotypes during clonal evolution. Alternatively, kar2 and kar3 in BPK081 cl8 may have originated independently from kar1 by simultaneous amplifications of multiple chromosomes. However, the presence of potentially triploid cells which resemble kar2 and kar3 opens other possibilities. On one hand, polyploidization has been demonstrated as an important mechanism in yeasts for quickly generating multiple and highly discrepant aneuploid karyotypes from a single parent through assorted mis-segregation of chromosomes during downstream cell divisions (Gerstein et al., 2015). In case a similar mechanism occurs in *Leishmania*, these 3n karyotypes found in BPK081 cl8 could represent an intermediate step between whole genome polyploidization event and reversion to aneuploid kar2 and kar3. On the other hand, 3n karyotypes could be reminiscent of hybridization, which was recently shown to occur in vitro (Louradour et al., 2020); the common observation of 3n karyotypes in *Leishmania* after hybridization in sand flies supports this hypothesis (Akopyants et al., 2009; Inbar et al., 2019, 2013; Romano et al., 2014).

Surrounding the 3 major karyotypes in the network of BPK081/0 cl8, other minor karyotypes with single somy alterations are observed, suggesting that once a successful karyotype expands, small variations of it are continuously generated by small changes in somies. This pattern is more evident in the karyotype network of BPK282/0 cl4, where almost all karyotypes which are found in at least 2 cells are at one somy change distance from another karyotype, suggesting that these karyotypes were also continuously generated by cumulative steps of small somy alterations. Accordingly, the founder karyotype of this clone (likely kar1) was already highly aneuploid and well adapted to culture, as the parent population from which BPK282/0 cl4 was isolated was already in culture for 21 passages (supp. fig. 1).

Highly aneuploidy Kps are observed in most in vitro cultured *Leishmania* promastigotes analysed so far by BGS (Franssen et al., 2020; Imamura et al., 2016; Van den Broeck et al., 2020). This usually affects a specific group of chromosomes, largely overlapping with the 8 polysomy-prone chromosomes described here. The early amplifications reproducibly observed in the Kp of parasite populations in transition from in vivo to in vitro (Domagalska et al., 2019; Giovanni Bussotti, a et al., 2018) suggest an adaptative role for specific polysomies in adaptation to culture. However, the mechanisms that determine which chromosomes are amplified are still poorly understood.

By investigating which chromosomes were more prone to somy alterations in rare and common karyotypes, we gathered evidence suggesting that all chromosomes can be stochastically amplified during population expansion, potentially at different rates, but selective forces likely dictate the higher frequency of polysomies observed in some chromosomes. Changes in the average chromosome copy numbers of cell populations are directly reflected in the average amount of transcripts encoded by the genes present on these chromosomes (Barja et al., 2017; Dumetz et al., 2017) and to a certain degree also affect the average amount of certain proteins (Cuypers, 2018). Consequently, aneuploidy might lead to dosage imbalances between the product of genes located in chromosomes that display different somies. The frequently observed co-modulation of multiple chromosomes – estimated with Pearson correlations here and across the Kp of 204 *L. donovani* isolates as previously described (Barja et al., 2017) – might reflect a dynamic compensation mechanism that reduces these imbalances and at the same time increases the dosage of key genes. Our GO analyses did not reveal any enrichment of biological functions in the (co-)amplified chromosomes. However, we observed an enrichment of snoRNA genes in some of the polysomy-prone chromosomes, accordingly Chr05, Chr26, Chr33 and Chr35. This class of small RNAs is involved in the extensive processing of ribosomal RNA (rRNA) characteristic of trypanosomatids, directly affecting ribosomal biosynthesis and ultimately translation, both increased in cultured promastigotes (Jara et al., 2017; Martínez-Calvillo et al., 2019). Amplification of these chromosomes as seen in many cells in vitro might ultimately boost the translation capacity of the cells due to a consequent higher abundance of snoRNAs. At the time of submission of the present article, a very recent study supporting and further addressing this hypothesis was pre-printed (Piel et al., 2021).

The high diversity of karyotypes identified in both models here described is in agreement with the idea of mosaic aneuploidy being a constitutive feature in *Leishmania* (Lachaud et al., 2014). The generation of karyotypic heterogeneity represents a source of functional diversity, due to variations in genes dosage (Dumetz et al., 2017), and it is also expected to facilitate the removal of detrimental mutations and the fixation of beneficial haplotypes (Barja et al., 2017; Sterkers et al., 2012). Although in a given environment some very different karyotypes might be limited to low frequencies, they may provide to the population a major (pre-)adaptation potential to unpredictable environmental changes, such as a change of host or drug pressure associated to chemotherapy (Dumetz et al., 2018, 2017; Shaw et al., 2016, 2020). Time-lapse SCGS studies of populations of parasites during clonal expansion under stable or varying environments are needed to test this pre-adaptation hypothesis. Combining SCGS with single-cell transcriptomics could also allow to understand better the impact of gene dosage imbalance on transcription with a single-cell resolution. Thus, high throughput single-cell sequencing methods represent a remarkable tool to understand key aspects of *Leishmania* biology and adaptability.

## Acknowledgements

We thank Prof. Dr. Thierry Backeljau for comments on the manuscript. This study received financial support from the Flemish Ministry of Science and Innovation (SOFI Grant MADLEI) and the Flemish Fund for Scientific Research (FWO, post-doctoral grant to FVdB). A.Y. was recipient of a grant from the Agence Nationale de la Recherche (ANR) within the frame of the “Investissements d’avenir” programme (ANR 11-LABX-0024-01 “ParaFrap”).

## Author contributions

All authors have approved the submitted version of this manuscript and have agreed both to be personally accountable for their own contributions and to ensure that questions related to the accuracy or integrity of any part of the work are appropriately investigated and resolved. This work was conceived and designed by GN, JCD & MAD. Data were acquired and analyzed by GN, PM, HI, IM, NK, AY, YS, JCD and MAD. Data interpretation was made by GN, PM, FVdB, YS, JCD and MAD. Paper was drafted by GN, PM, JCD and MAD and substantively revised by HI, FVdB and YS.

## Supplementary text

### Definitions and Glossary

In the present paper, we use the following definitions for population, strains and clones; adapted from the nomenclature of salivarian trypanosomes (Baker et al., 1978). Accordingly:

- A population is a group of *Leishmania* cells present at a given time in a given culture or host;
- A strain is a population derived by serial passage in vitro from a primary isolate (in our case, from patient samples) without any implication of homogeneity but with some degree of characterization (in our case bulk genome sequencing).
- A clone is derived from a strain and is a population of cells derived from a single individual presumably by binary fission.

Other terms are defined in the following glossary:

| Term | Definition |
| --- | --- |
| Bulk Genome Sequencing (BGS) | Whole genome sequencing performed in a group of cells combined as a single sample. |
| Single Cell Genome Sequencing (SCGS) | Genome sequencing performed in single cells individually. |
| Somy | The number of copies of a given chromosome in a cell. |
| Polysomy | A somy higher than 2. |
| Karyotype | The set of copy numbers of all chromosomes in a cell. |
| Cell Karyotype | The karyotype of a cell determined by SCGS. |
| Populational Karyotype (Kp) | The average karyotype of a population determined by BGS. |
| Ploidy | The most frequent somy in a karyotype. |
| Euploidy | A condition where all chromosomes display the same somy in a cell. |
| Aneuploidy | A condition where one or more chromosomes display a somy that diverges from the other chromosomes in the same cell. |
| Mosaic Aneuploidy | A condition where different aneuploid karyotypes co-exist in the same population. |
| Cell scale factor (Sc) | The lowest number between 1.8 and 5 by which when the average normalized read depths of all chromosomes in a cell are multiplied the resulting numbers are the closest to integers as possible. |
| Raw somy | The average normalized read depth of a chromosome multiplied by Sc. |
| Integer somy | The integer value assigned to a raw somy. |

### Supplementary methods

#### Single-cell DNA sequence data analysis

Illumina Base call files (BCL) were demultiplexed and converted to FASTQ files using the cellranger-dna mkfastq command of the CellRanger^TM^ DNA pipeline (10X Genomics). The FASTQ files were then used as inputs to the cellranger-dna cnv command in order to associate reads to individual cells based on their 10X barcodes and to map reads to a customized version of the LdBPKv2 *L. donovani* reference genome (available at ftp://ftp.sanger.ac.uk/pub/project/pathogens/Leishmania/donovani/LdBPKPAC2016beta/), where ‘N’s were added to the ends of chromosomes 1 to 5 to reach the 500kb minimum size allowed by the CellRanger DNA pipeline. The pipeline divides the genome into adjacent 20kb bins and outputs a CSV file containing the number of reads mapped to each bin. This file was used to estimate chromosomes copy number in a custom script written in R.

An overview of the steps performed by the script is shown in supp. fig. 3A. The script first removes bins with a low number of mapped reads by eliminating any bin showing an average depth of 0.5 read/cell. Then, the difference between the median number of reads of each bin and the chromosomal median is calculated. Bins with outlier values are determined using the boxplot.stats function from the R package grDevices v3.6.2. These outlier bins are removed from downstream analysis (supp. fig. 3B). This also excludes common local-CNVs found in some *donovani* strains, as for instance the H-Locus and the M-Locus in Chr23 and Chr36 respectively (Downing et al., 2011), present in the BPK strains/clones but absent in the HU3 strain. After removal of outlier bins, the bins depths are normalized by the cell mean and are used to estimate intrachromosomal variation (ICV). ICV is determined for each cell by dividing each chromosome in 3 segments and calculating the ratio between the segment with the highest and the segment with lowest depth. The mean of the five highest ICV values (i.e. the 5 most variable chromosomes in a cell) is assigned as its ICV-score. The distribution of ICV-scores in each sample was graphically analyzed in order to determine a threshold for exclusion of noisy cells. This threshold was defined as 2.0 for BPK282 cl4 and 1.7 for the BPK081 cl8 and the ‘super-mosaic’ samples.

The copy number of chromosomes in a cell is defined based on their normalized mean depth (NMD), i.e., the mean of the normalized depth values of the 20kb bins of a chromosome. In this sense, NMDs reflects the relative differences in copy number between chromosomes, but absolute copy numbers must be inferred based on the ratios between NMDs of different chromosomes in a cell. Thus, considering that chromosomes copy numbers must be integers, the script uses an approach to determine absolute copy numbers which consists of multiplying NMDs by a scale factor which minimizes distances between the multiplied NMDs and integers. Therefore, the scale factor is defined as the lowest value between 1.8 and 5 which results to the closest approximation of NMDs to integers when they are multiplied by this factor. As the scale factor is directly affected by the ploidy of the cell, the limitation of the scale factor to values higher than 1.8 heuristically assumes that the lowest baseline ploidy of a cell is 2n. This was done to prevent that 2n cells with no odd somy value would be scaled as 1n cells.

In order to determine the scale factor, the script multiply the NMDs of a cell by 1000 equidistant numbers between 1.8 and 5. For each multiplication, the difference between the resultant values and their closest integers is calculated for each chromosome and averaged. The value that results in the lowest average distance to integers is then assigned to the cell as its scale factor (supp. fig. 3C). In case two or more scale factors result in the same average distance to integers, the one with the lowest value is chosen.

Since *Leishmania* chromosomes are biased in GC content (Imamura et al., 2020), with small chromosomes (Chr1 to Chr5) displaying a higher GC content than others, amplification bias due to differences in GC content can have a negative impact in the determination of the copy number of these chromosomes. Plotting the distribution of NMD values leads to different peaks, each peak representing one of the somy values, however, the peaks of these small chromosomes with high GC content are shifted relative to the other chromosomes (supp. fig. 3D upper panel). Thus, to compensate for chromosome-specific amplification and to further define the somies of the cells, the above explained scale factor are used at two levels, i.e., at population level (all cells combined) as well as at single cell level (defined for each cell). In this sense, the script first defines a single scale factor to the whole population (Sp) by which NMDs are multiplied and the distribution peak of the scaled NMDs of each chromosome is adjusted to the closest integer (supp. fig. 3D bottom panel). Then, these values are divided back by Sp and based on this output a second scale factor is defined for each cell (Sc). Thus, the NMDs of the chromosomes in a cell after bias compensation multiplied by the cell’s Sc defines the ‘raw somies’ of the chromosomes of that cell.

Despite the fact that the abovementioned steps have moved the NMDs distribution closer to integer values, those values are still floating-point numbers. To determine the cells karyotypes, the raw floating-point somies are converted to integer copy numbers using Gaussian Mixture Models (GMMs). To generate a GMM for each chromosome, a vector containing all raw somy values determined for that chromosome among the filtered cells in a sample is used as input to the normalMixEM function of the mixtools R (Benaglia et al., 2009), following the defined rules bellow:

1) The possible integer values are defined as the number of different integers found when all values in the vector are rounded to the closest integer.
2) The number of components (k) is determined as the total number of possible integer values.
3) The ratio between means (µ) of k gaussians are constrained to the ratios between the possible integer values.
4) If for a given gaussian, less than 5% of the values are inside the interval between µ-0.2 < µ < µ+0.2, the standard deviation (σ) of that gaussian is arbitrarily limited to 0.1.
5) At least 5 iterations must be performed before a gaussian is defined.

Thus, for each chromosome in a sample, a gaussian is built for each possible integer somy (supp. fig. 3E). Raw somies are then converted to the rounded µ of the gaussian of which they have the higher probability of belonging to. Since the GMMs must be built between cells sharing the sample baseline ploidy, and as the vast majority of cells in all samples sequenced in the present study had a scale factor lower than 2.5 and consequently were considered 2n cells (supp. fig. 3F), the GMMs were applied only to 2n cells. Moreover, since the number of non-2n cells were always very low, GMMs could not be built separately for cells with other baseline ploidies. Thus, cells which baseline ploidy was different than 2 were treated differently. In this case, cells with intermediate somies, i.e, with at least one raw somy values that are at a distance greater than 0.4 from its closest integers, were considered unresolvable and were removed from downstream analysis. The reminiscent had their raw somy values simply rounded to the closest integer. Karyotypes were then defined as the concatenated set of integer somy values found in a cell.

#### Doublet detection

Two different methods were used for doublet detection, i.e. an in-house developed methodology and Demuxlet (Kang et al., 2018), both exploiting the difference in SNP profile between HU3 cells versus other cell lines.

The in-house developed approach uses the following methodology: 1) Homozygote SNPs for the HU3 strain are predicted based on the genome of the HU3 strain sequenced by BGS (data not shown). 2) For each of those HU3 homozygote SNPs, the occurrence of this SNP is derived for each of the single cells in the 4-strains mixture sample (further called ‘super-mosaic’). Given the low sequencing depth per cell (on average around 1x), this will report the absence or presence for each SNP. 3) For the HU3 cells in the super-mosaic, the majority of SNPs should be detected, while for the other three strains no SNPs should be detected. In case of a doublet consisting of a HU3 cell with a cell from one of the other three strains, two different scenarios can occur: If the sequencing depth is low, only the allele of one of the two cells can be predicted, while in case of a sufficient sequencing depth (at least 2x), both alleles (either the HU3 or the reference allele) can be detected, resulting in an allele frequency of 50%. In both cases, overall detection rate of the homozygote SNPs should be around 50%. In order to compensate for sequencing errors and differences in sequencing depth, libraries detecting between 10% and 90% of the HU3 homozygote SNP list were classified as doublets. Homozygote SNPs where predicted based on the genome of the HU3 strain. Genetic variants were detected using the mpileup and call command of BCFftools (version 1.10.2). The view and query command of BCFtools were used to filter out genetic variants fulfilling the following conditions 1) minimum sequencing depth of 100, 2) only SNPs i.e., removing indels, 3) biallelic, 4) homozygous. In a second step, for each single cell those SNP positions are checked using the bcftools mpileup command.

Demuxlet was run using the default parameters with the following input: 1) the bam file returned by the Cell Ranger software, produced for the single-cell experiment with the super-mosaic, 2) a vcf file describing the two different SNP profiles, i.e., the SNP profile for HU3, and the SNP profile for the three other strains.

### Supplementary results & discussion

#### Sequencing statistics

Summary of sequencing statistics is provided in table S1. The BPK282 cl4 and BPK081 cl8 were sequenced with the same targeted depth (75.000 reads per cell) but BPK282 cl4 sample displayed a depth which was lower than anticipated (29.192 reads per cell). This was due to a high fraction (53.3%) of reads without a cell barcode in this sample, which according to the manufacturer indicates free floating DNA or a problem during library prep, but which unlikely affect copy number estimation. The scCNV library of the super-mosaic sample was sequenced deeper (209.000 reads per cell) to better allow the distinction between doublets. Higher coverage depths per cell were also associated with lower intra-chromosomal variation and lower frequency of intermediate somy values (supp. fig. 4A-B). This explains why sample BPK282 cl4 displayed a higher overall ICV score compared to the other samples.

The noisy nature of whole genome amplification ultimately leads, in some cases, to the existence of raw somy values are at similar distances from two integers. Although the conversion of raw somies into integers could be achieved by simply rounding the raw somy values to the closest integers, this could overestimate the number of karyotypes identified in a population, as the wrong determination of a somy value of a single chromosome in a single cell is sufficient to lead to a new artificial karyotype. Thus, in order to convert the raw somy values into integers, we used a more stringent approach by constructing GMMs based on the distribution of raw somy values of each chromosome among cells in a given sample. One of the consequences of using this approach is that the frequency of which an integer somy value is present in a population influences the probability of a raw somy value to be assigned to this integer. This favors that intermediate somy values are assigned to the most frequent integer somy values in the population, reducing the chances of misinterpreting an intermediate value as a new, rare integer, and consequently greatly reducing the number of artificial karyotypes caused by the misinterpretation of a somy. This is evident, for example, when comparing the number of karyotypes identified in the BPK282 cl4 sample using the GMMs (207 karyotypes) and when raw somies are just rounded to their closest integers (525 karyotypes - supp. fig. 4D).

Noisy data had also an impact on the scaling of the NMDs of cells into raw somies, as differences between chromosomes NMDs becomes less discrete. In the 3 samples submitted to SCGS here we noticed a higher ICV-score in a large fraction of cells which were scaled to baseline ploidies different than 2 (supp. fig. 4C). These cells were removed from karyotype estimation either due to their ICR-score being above the threshold, or due to the presence of unresolvable intermediate somy values as described in the supplementary materials and methods.

## Supplementary Figures

**Supplementary figure 1.**
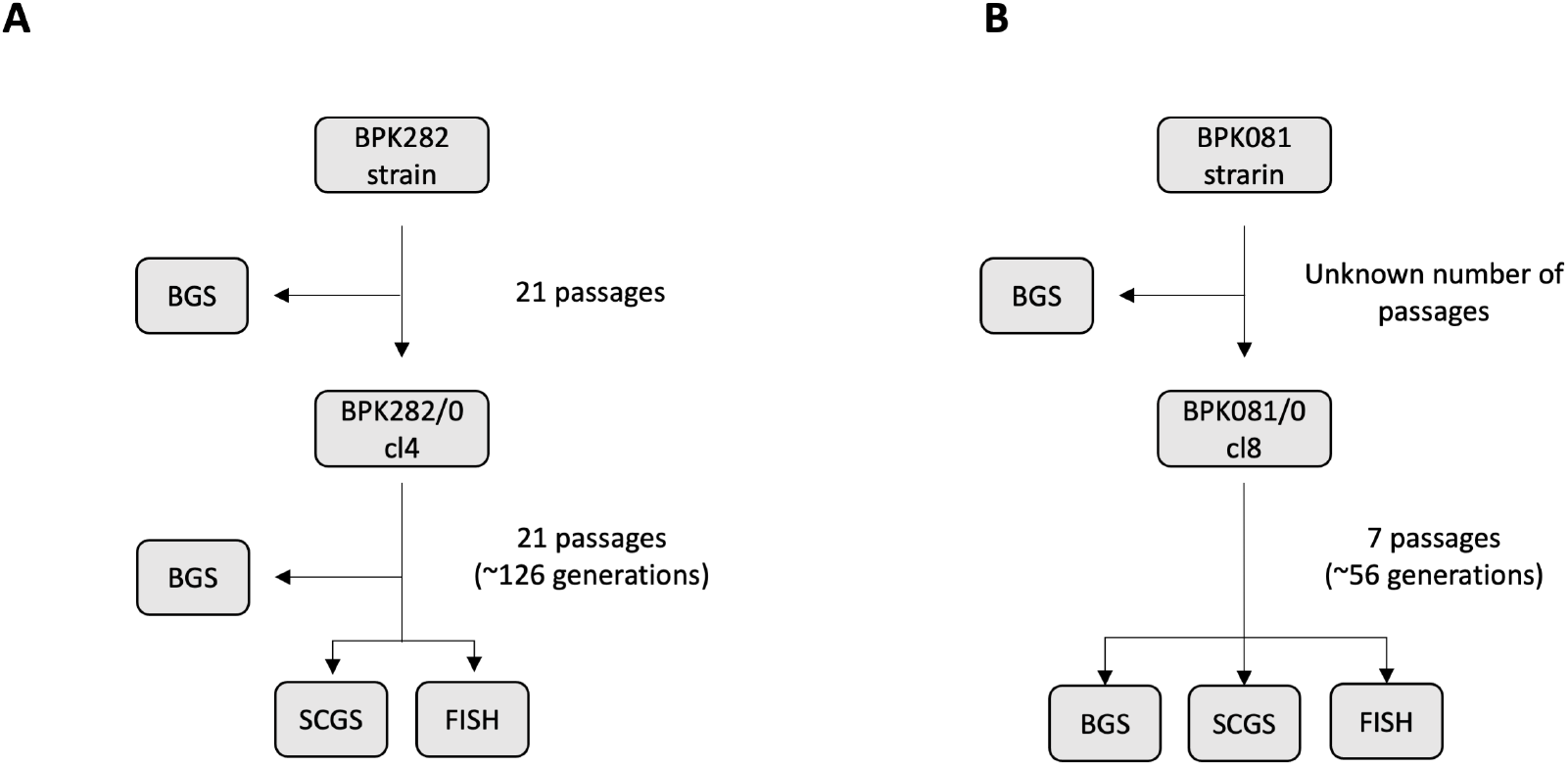
Flow chart of the two clonal populations used in the present study. For BPK282 cl4, SCGS and FISH were performed in cultures at the same passage number. BGS was performed previously at passage 13 after cloning. For BPK081 cl8, all experiments were performed with the same culture. Number of generations is roughly estimated as 26+((p-1)*5)), where p is the number of passages. This is done assuming that it takes about 26 generations to reach a total of ∼7×10^7^ cells starting from 1 cell, an approximation to the total number of cells usually found in a culture flask with 5mL of culture medium at the moment the first passage is done, and also assuming that each subsequent passage represents ∼5 generations.

**Supplementary figure 2.**
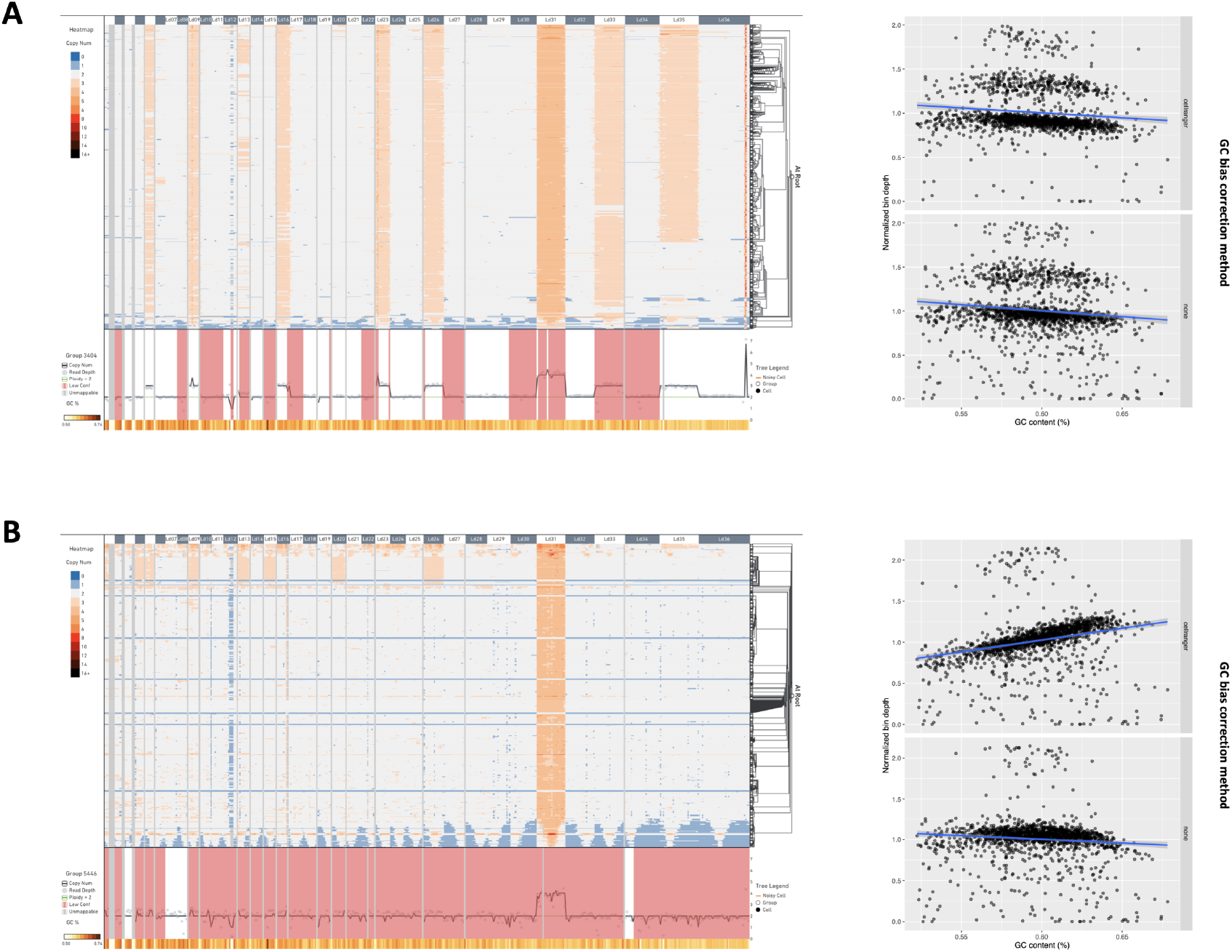
CNV profile of BPK282 cl4 and BPK081 cl8 calculated with the Cell Ranger^TM^ pipeline and visualized with the Loupe^TM^ scDNA Browser software (10X Genomics). In each sample, cells (rows) are arranged in 512 clusters, the maximum number of clusters allowed by the software. CNVs (columns) are depicted in windows of 80kb. Insets on the left display the effect of the the GC bias correction algorithm of the Cell Ranger^TM^ pipeline on the normalized read depth of bins (top) when compared to no bias correction (bottom).

**Supplementary figure 3.**
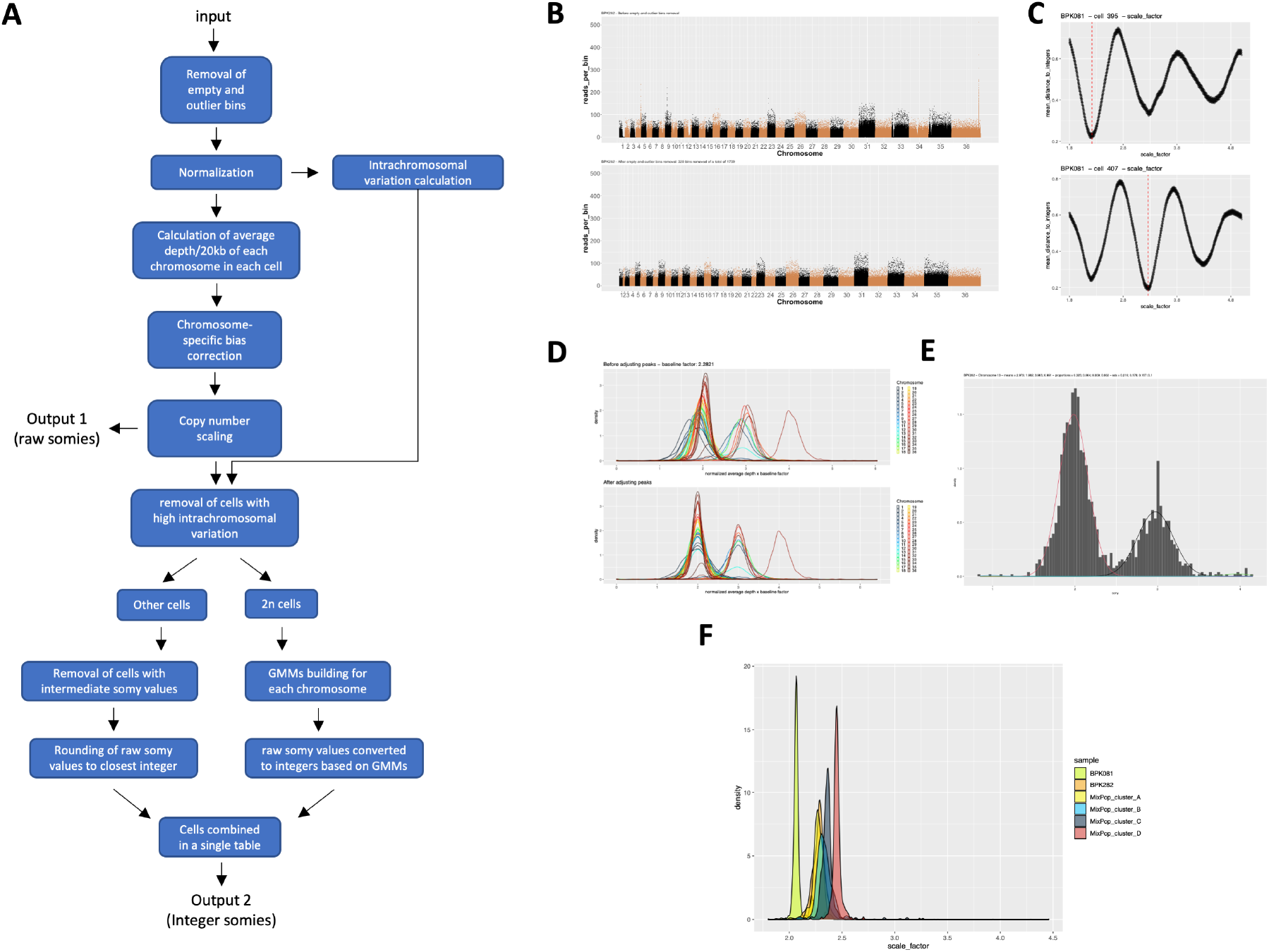
Bioinformatics pipeline for somy estimation. **A.** Flow chart of the script developed to estimate chromosomes copy numbers based on their average depth/20kb bin. The input file is a matrix containing the read count of each 20kb bin for each cell. Two output files are generated, one with the raw somy values (floating points) and another with integer somy values. **B.** An example of the effect of the removal of empty and outlier bins in the BPK282 cl4 data. In this step, small intrachromosomal CNVs are also removed. **C.** Example of the determination of the scale factor for a 2N cell in the BPK081/0 cl8 sample with karyotype 2 (top panel) and a 3N cell with karyotype 13 (bottom panel). Y-axis represents the mean distance to integers when the NMDs of that cell are multiplied by a given scale_factor (x-axis). Red dashed line denotes the scale factor value defined for that cell. **D.** An example of the chromosome-bias correction step in the BPK282 cl4 data. **E.** Example of a Gaussian Mixture Model (GMM) built for chromosome 13 in the BPK282 cl4 data. The histogram represents the distribution of raw somy values for this chromosome in this sample, while the gaussian curves represent the GMM built for it. In this step, a gaussian is built for each integer, and raw somy values are assigned to the integer corresponding to the gaussian to which they have the higher probability. **F.** Distribution of the scale factors between all cells sequenced in this study.

**Supplementary figure 4.**
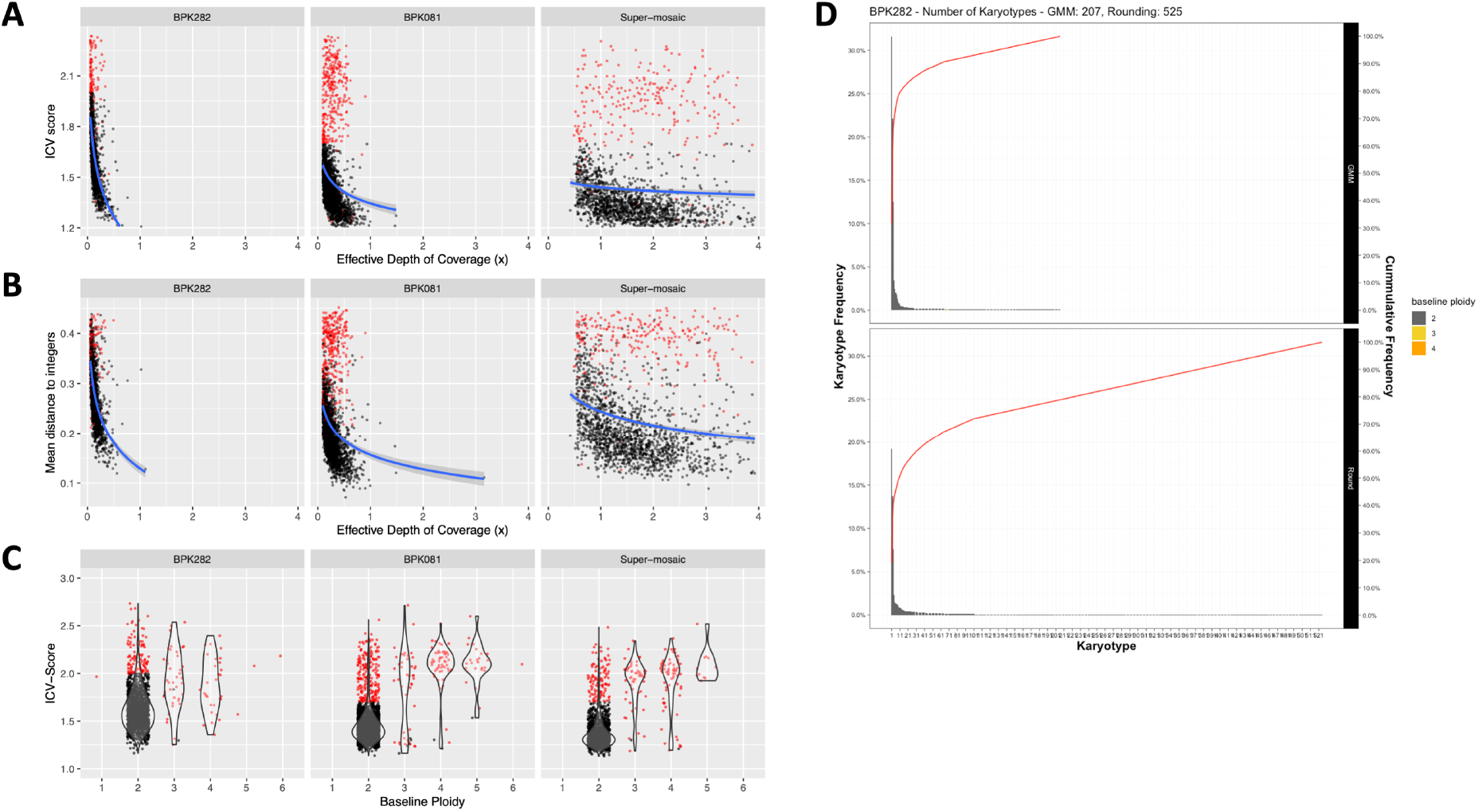
The relationship between the depth of coverage per cell and the cells ICV-score (**A**), mean distance to integers (**B**) and the relationship between the baseline ploidy defined for a cell – which is a direct consequence of the cells scale factor – and the cells ICV-score (**C**). Red dots represent cells which were removed from karyotype estimation. **D.** Comparison of the number and distribution of karyotypes identified in BPK282 when using the GMMs (top) and when raw somies are simply rounded to their closest integers (bottom).

**Supplementary figure 5.**
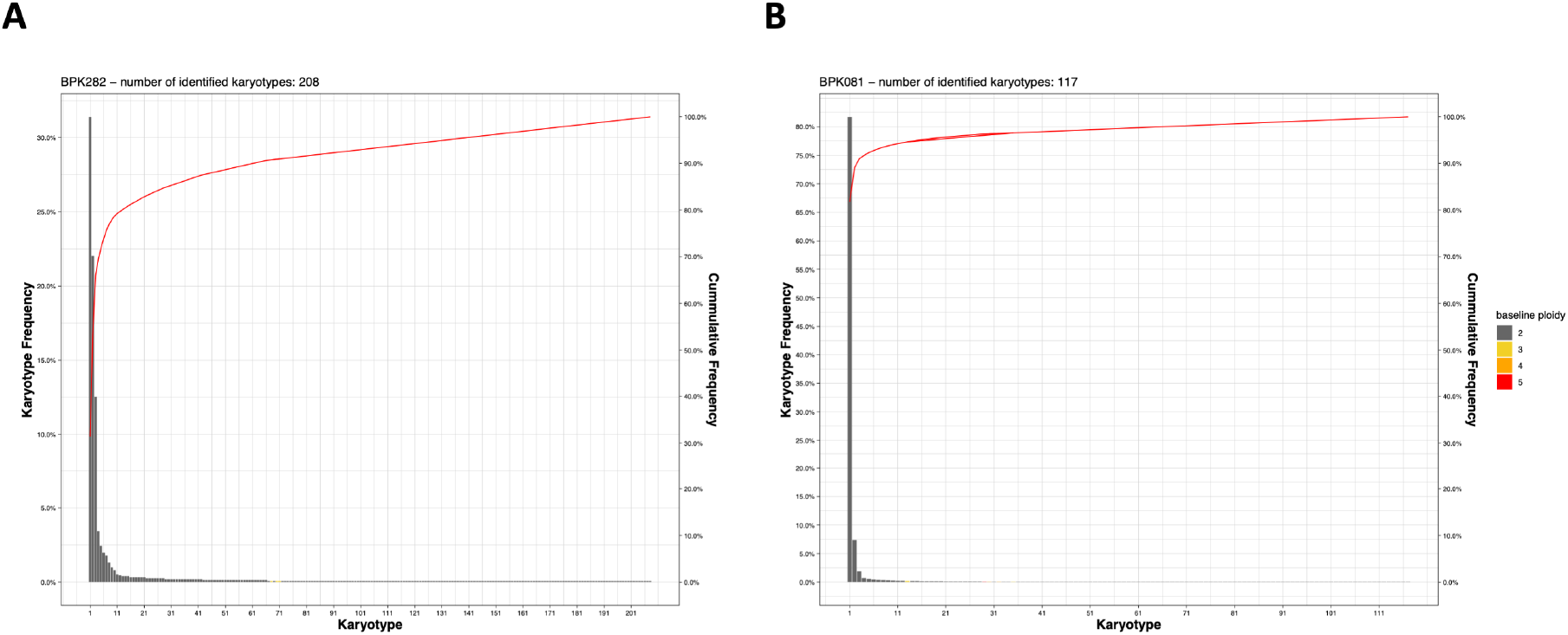
Frequency distribution of the karyotypes identified in **A.** BPK282 cl4 and **B.** BPK081 cl8 clones.

**Supplementary figure 6.**
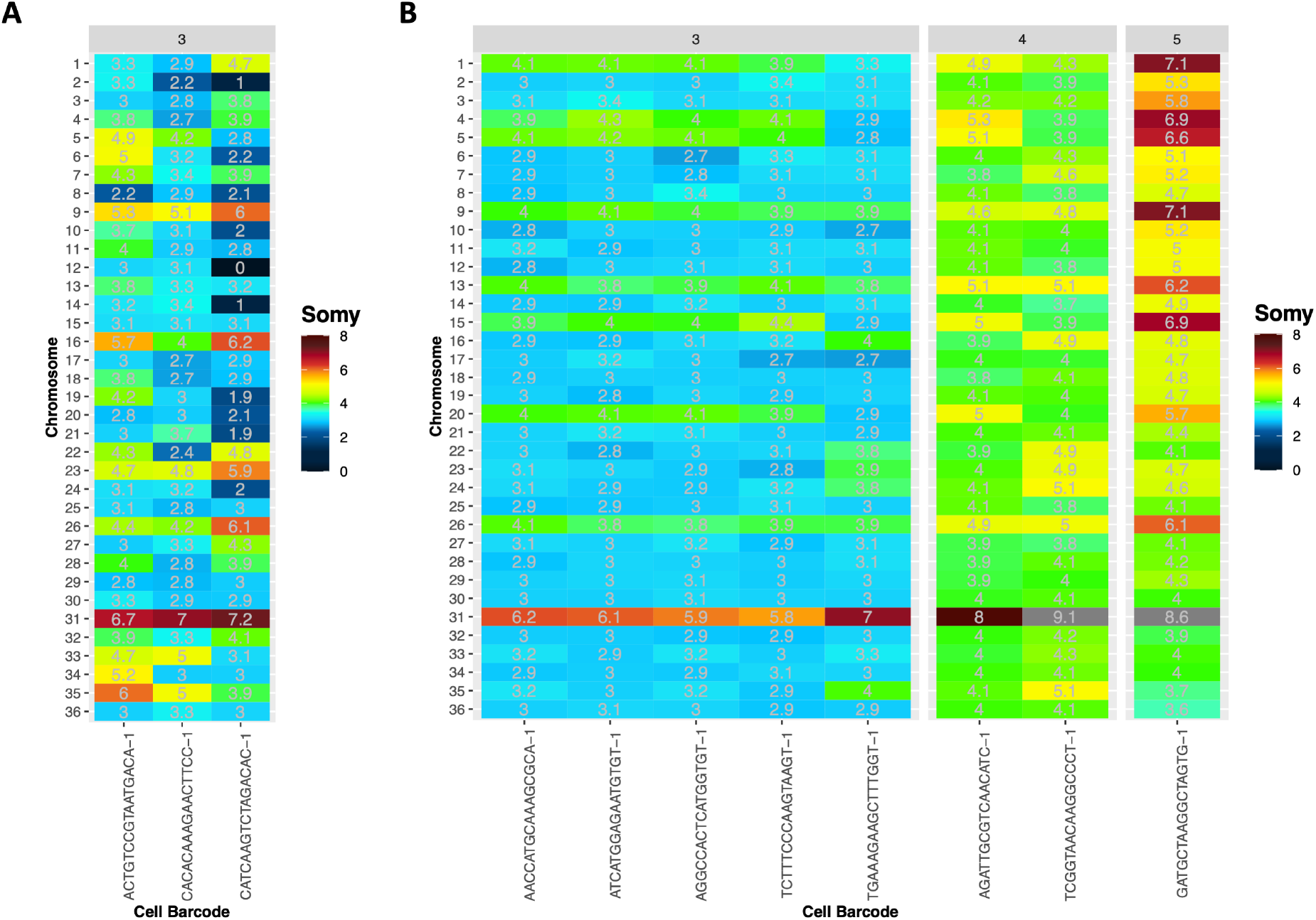
Raw somy values of potentially polyploid cells in BPK282 cl4 (**A**) and BPK081 cl8 (**B**) clones. Plots are separated by the baseline ploidy of the cells (indicated in the top).

**Supplementary figure 7.**
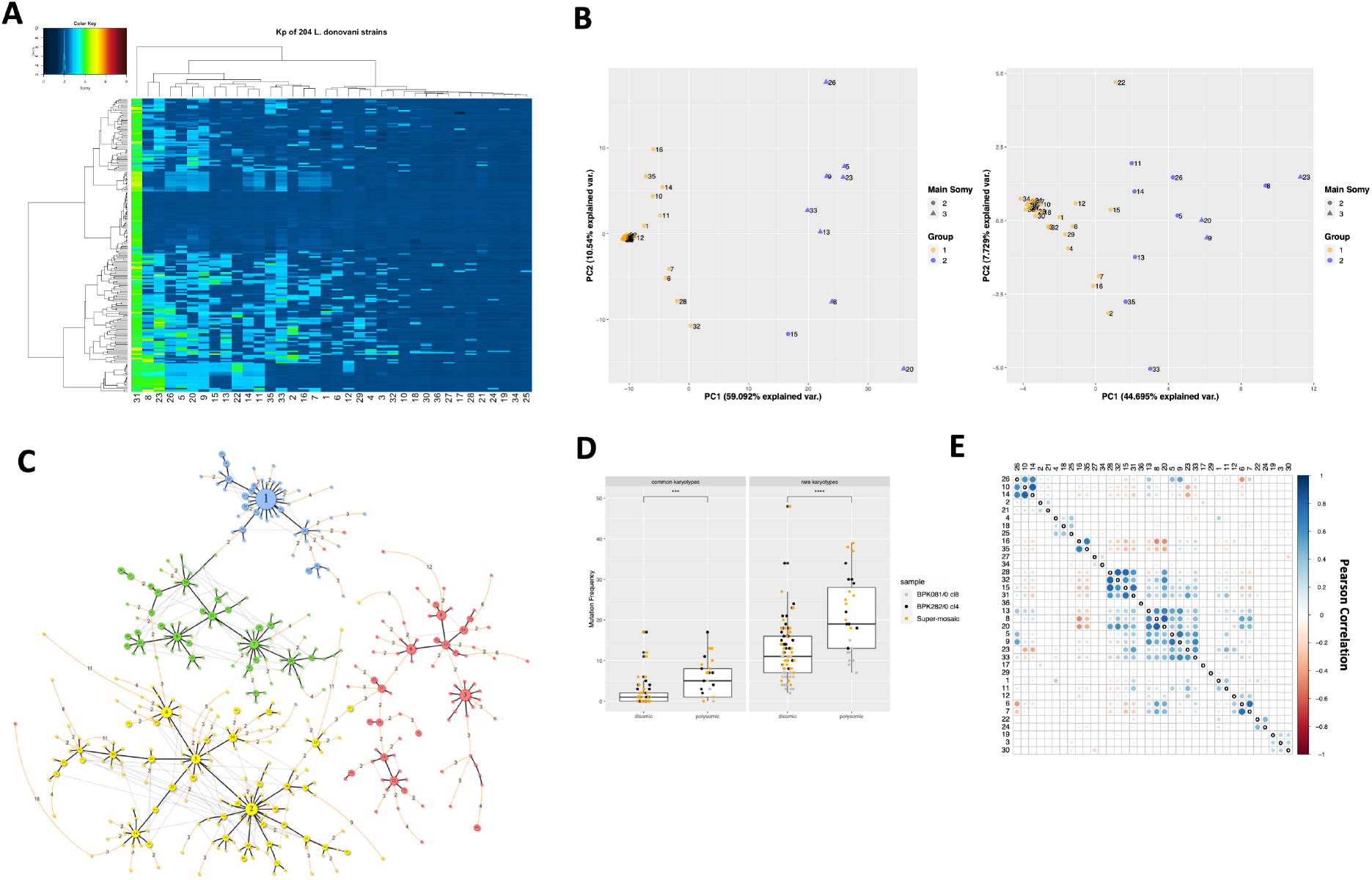
Supporting images for Fig 3 in the main text. **A.** Somies observed in the BGS data of 204 *L.donovani* strains (rows) with chromosomes (columns) hierarchically clustered. Data from Imamura H. et al, 2016. **B.** Principal Component Analysis constructed based on the somy values of each chromosome (dots) found in the 1554 cells from 6 different strains/clones (left panel) or among the Kp’s of 204 *L. donovani* strains (right panel). **C.** Karyotype Network of the ‘super-mosaic’ population. Color of the nodes indicate the cluster to which each karyotype belongs. Yellow: Cluster A; Red: Cluster B; Green: Cluster C; Blue: Cluster D. **D.** Comparison of the frequency of somy change events between the polysomy-prone chromosomes and the chromosomes which are usually found as disomic. *** = p.value <0.001 and **** = p.value <0.0001 (T-test). **E.** Pearson correlation matrix used to generate the chord diagram in figure 3C in the main text. Correlations with p-value higher than 0.05 are not shown.

**Supplementary figure 8.**
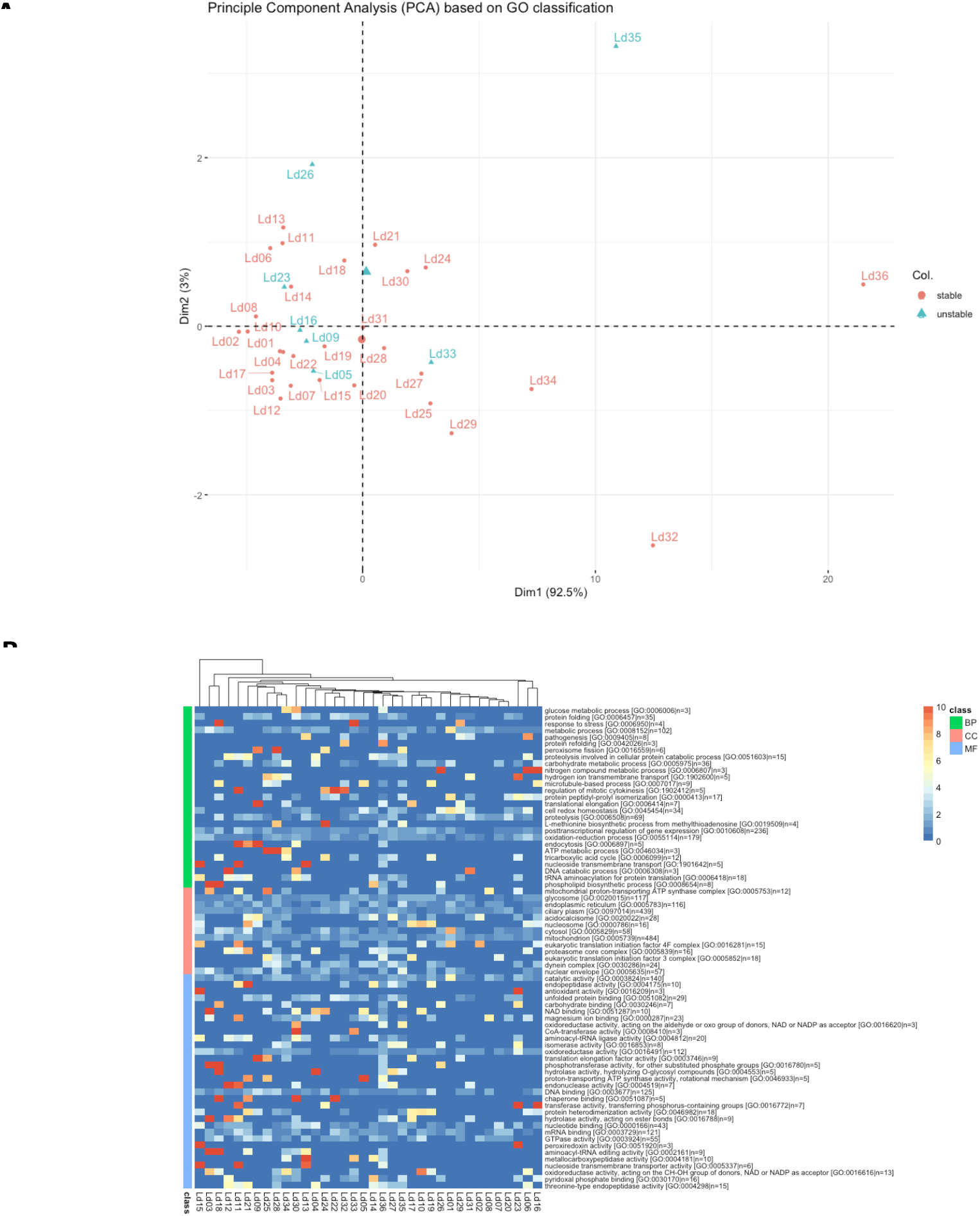
**A.** Principal component analysis (PCA) based on the Gene Ontology (GO) annotation. Based on the GO annotation provided by TriTrypDB, the percentage of each GO category (minimal category size set to 10, maximum category size set to 500), the chromosome by GO category percentage matrix is used as input for the PCA analysis. Chromosomes indicated as “stable” due to their stable disomy are indicated in red, chromosomes which showed frequent changes in ploidy level are indicated in cyan. No obvious clustering of unstable chromosomes is observed based on their GO classification **B.** Heatmap showing the ratio of the genes assigned to a GO class over the total number of genes per GO class (colour code between 0% and 10%), calculated per chromosome. The list of GO classes shown in this heatmap are significantly enriched promastigote-specific GO classes, derived based on the transcriptomics data as available in Dumetz et al. 2017, and are grouped over the three main categories i.e. Biologial Process (BP), Cellular Compartment (CC) and Molecular Function (MF). No clear clustering of polysomy-prone chromosomes based on the GO classification was observed.

